# Disconnectome Associated with Progressive Ischemic Periventricular White Matter Lesions

**DOI:** 10.1101/2019.12.24.888081

**Authors:** Zhengjun Li, Sudipto Dolui, Mohamad Habes, Danielle S. Bassett, David Wolk, John A. Detre

**Author notes:** **Corresponding author:** John A. Detre, M.D., **Email:**, Department of Neurology, University of Pennsylvania, 3W Gates Pavilion, 3400 Spruce Street, Philadelphia, PA 19104, USA. **Author Contributions:** Designed research: ZL, DB, DW, JD Performed research: ZL, JD Contributed tools: SD, DB, DW Analyzed data: ZL Wrote the paper: ZL, SD, MH, DB, DW, JD.

## Abstract

Periventricular white matter (PVWM) hyperintensities on T2-weighted MRI are ubiquitous in older adults and associated with dementia. Efforts to determine how PVWM lesions impact structural connectivity to impinge on brain function remain challenging in part because white matter tractography algorithms for diffusion tensor imaging (DTI) may lose fidelity in the presence of lesions. We used a “virtual lesion” approach to characterize the “disconnectome” associated with periventricular white matter (PVWM) lesions. We simulated progressive ischemic PVWM lesions using sub-threshold cerebral blood flow (CBF) masks derived from a previously published group-averaged map acquired from N=436 middle aged subjects in which the lowest CBF values were seen in PVWM and morphologically recapitulated the spatial pattern of PVWM hyperintensities seen in typical aging. We mimicked the age-dependent evolution of PVWM lesion burden by varying the threshold applied to the CBF map. We found that the optic radiations, inferior fronto-occipital fasciculus, inferior longitudinal fasciculus, corpus callosum, temporopontine tract and fornix were affected in early simulated PVWM lesion burdens, and that the connectivity of subcortical, cerebellar, and visual regions were significantly disrupted with increasing simulated PVWM lesion burdens. We also validated the use of virtual lesions to simulate the disconnectome due to WM hyperintensities in a cognitively normal elderly cohort (N=46) by evaluating correlations between structural and functional connectomes. The virtual lesion approach provides new insights into the spatial-temporal changes of the brain structural connectome under progressive PVWM burdens during normal aging.

**Significance Statement:** We determined the disconnectomes caused by periventricular white matter (PVWM) lesions using the “virtual lesion” approach. We validated the approach using lesions, DTI and resting-state fMRI data from elderly subjects. We simulated disconnectome of progressive PVWM lesions using cerebral blood flow (CBF) masks in PVWM region with normative DTI data, which provides specificity for an ischemic mechanism and begins to address the possibility that connectivity may be affected by reduced CBF prior to the development of overt lesions on T2-weighted FLAIR MRI. The current study presented new insights into the spatial-temporal evolutions of the brain structural connectome under progressive PVWM burdens under normal aging.

## Introduction

White matter (WM) lesions are typically detected as hyperintensities on T2-weighted fluid attenuated inversion recovery (FLAIR) sequence in magnetic resonance imaging (MRI), and are highly prevalent in older adults (1), including asymptomatic individuals (2). WM lesions in aging populations appear first and more frequently in the periventricular white matter (PVWM) (3) as variable size caps on the frontal and occipital horns, and as thin rims along the walls of the lateral ventricles on transverse sections of MRI scans, but with increasing severity may become confluent with lesions that occur in the deep white matter (DWM). PVWM lesions are more ubiquitous than DWM lesions (4, 5), and may be specifically associated with cognitive and functional decline (6–8).

PVWM lesions are thought to result primarily from ischemic microangiopathy (1), but are not specific for this mechanism. Postmortem correlations with MRI demonstrate that PVWM lesions are associated with demyelination and axonal loss (9), although histopathological findings are also heterogeneous (10). PVWM lesions, along with DWM lesions, lacunar infarcts, and microhemorrhages, underlie the syndrome of subcortical vascular cognitive impairment and also contribute to dementia accompanying Alzheimer’s disease (1,11–14). However, the mechanisms by which PVWM lesions selectively alter cognitive function remain uncertain, in part because these lesions are typically seen with comorbid brain pathologies.

White matter lesions classically cause disconnection syndromes by interrupting communication between the brain regions that they connect, such as hemiparesis with lesions affecting the corticospinal tracts. The distributed network properties of the brain can be studied noninvasively using diffusion tractography to infer structural connectivity in WM or using correlated fluctuations in BOLD fMRI to study functional connectivity between gray matter regions (15). Prior research using DTI data acquired in patients with small vessel disease has demonstrated alterations in structural network properties that correlate with cognitive decline (16–19). Yet, to date the specific connectivity of the periventricular white matter has not been elucidated. Given the ubiquity of PVWM hyperintensities even in healthy aging, identifying the pathways affected is of interest in further characterizing the mechanisms and progression of age-associated cognitive decline. Although early stage PVWM hyperintensities are small, their impact on structural connectivity may be relatively diffuse since multiple white matter tracts cross as they skirt around the ventricles. Furthermore, PVWM hyperintensities have been shown to have an ischemic penumbra (20–22), and it is quite possible that reduced CBF may result in impaired WM connectivity even prior to the appearance of overt hyperintensities.

Two main factors make it challenging to determine the impact of PVWM lesion on structural connectivity in the adult human brain. Firstly, lesions can alter the diffusion properties of WM and disrupt computerized tract-tracing algorithms, resulting in abnormal structural connectomes (23–27). Secondly, while the PVWM shows by far the highest WM lesion frequency across cohorts, individual subjects with PVWM lesions typically also have lesions elsewhere in the WM. To circumvent these limitations, in this study, we used a “virtual lesion” method to conduct DTI fiber tracking using young healthy subjects’ diffusion data from the Human Connectome Project (HCP) along with simulated PVWM lesions inserted as a region of avoidance (ROA) for the tractography algorithm.

To simulate PVWM lesions, we built upon our recent work demonstrating that the distribution of PVWM hyperintensities corresponds to the distribution of white matter perfusion, which is lowest in the periventricular region (28). In our prior study, we characterized spatial variations in white matter CBF using a group CBF map obtained from a large cohort (N = 436) of middle-aged adults from the Coronary Artery Risk Development in Young Adults (CARDIA) study. We constructed masks simulating PVWM lesions by thresholding the group-averaged perfusion map. By varying the threshold of this group map, we are able to generate lesion masks that closely recapitulate the age-dependent evolution of PVWM lesion burden without contamination by lesions in other regions of white matter. These CBF-derived lesion maps were then used as ROA to selectively identify the WM pathways most likely to be affected by progressive PVWM disease. Although virtual WM lesions could also be simulated by thresholding WM lesion frequency maps, the use of thresholded CBF provides a more mechanistically specific link to small vessel ischemic disease.

To further support the notion that a virtual lesion approach can be used to infer the effects of WM lesions on structural connectivity, we also examined structure-function relationships in multimodal MRI data acquired in a cohort of 46 cognitively normal and amyloid-negative elderly subjects that included T2-weighed FLAIR images used to segment WM hyperintensities. We used these data in several ways to validate the use of thresholded group CBF data as a proxy for WM lesions. We first demonstrated a concordance between the distribution of PVWM hyperintensities in this cohort and the PVWM CBF gradient used for CBF-based lesion simulation. We then demonstrated a concordance between the virtual disconnectomes generated using a mask based on the WM lesions derived from the elderly cohort and a volume-matched mask based on CBF. Finally, building on the notion that structural and functional connectivity may be expected to correlate (29–32), we used resting-state fMRI data acquired from the elderly cohort to compare correlations between functional connectivity and structural connectivity measured using each subject’s DTI data, with and without using each subject’s WM lesion mask as a ROA, to demonstrate that structure-function correlations were improved after adding the ROA. Finally, we validated the use of normative HCP data by comparing each subject’s functional connectivity to their structural connectivity determined by adding their WM lesion mask to HCP data and showing that structure-function correlations were improved further. These validations demonstrate that progressive PVWM lesions can be simulated using thresholded CBF data and that performing tractography using high-quality HCP DTI data using lesion-based ROA can identify meaningful changes in structural connectivity.

## Results

### White matter lesion frequency versus periventricular cerebral blood flow

A WM lesion frequency map derived from the elderly subjects (N = 46, age range 58 to 88 yrs, mean ± std = 72.30 ± 6.81 yrs) is shown in Fig. 1A. WM lesion volumes ranged 0 – 14288 mm^3^, with median/mean/standard deviation = 668/1937/3033 mm^3^. Lesion frequency was highest in the PVWM; however, numerous lesions were also present in the DWM. Fig. 1B shows a map of cerebral blood flow (CBF) in standard space derived from arterial spin labeled perfusion MRI data in 436 middle aged subjects, as described in our prior work (28). This map has been overlaid onto the same slices shown in Fig. 1A and thresholded to only display voxels with group-averaged CBF in the 0-20 ml/100g/min range. As noted in our prior work, the distribution of these lowest perfused voxels in the brain closely recapitulates the pattern of PVWM lesions. Based on this correspondence, a pseudo-PVWM lesion mask can be created by thresholding the group CBF data at various levels to approximate the evolution of PVWM lesions without contamination from DWM lesions. Note that the CBF ≤ 20 ml/100g/min (entire region of color overlay) would correspond to essentially the entire PVWM being lesioned.

**Figure 1.**
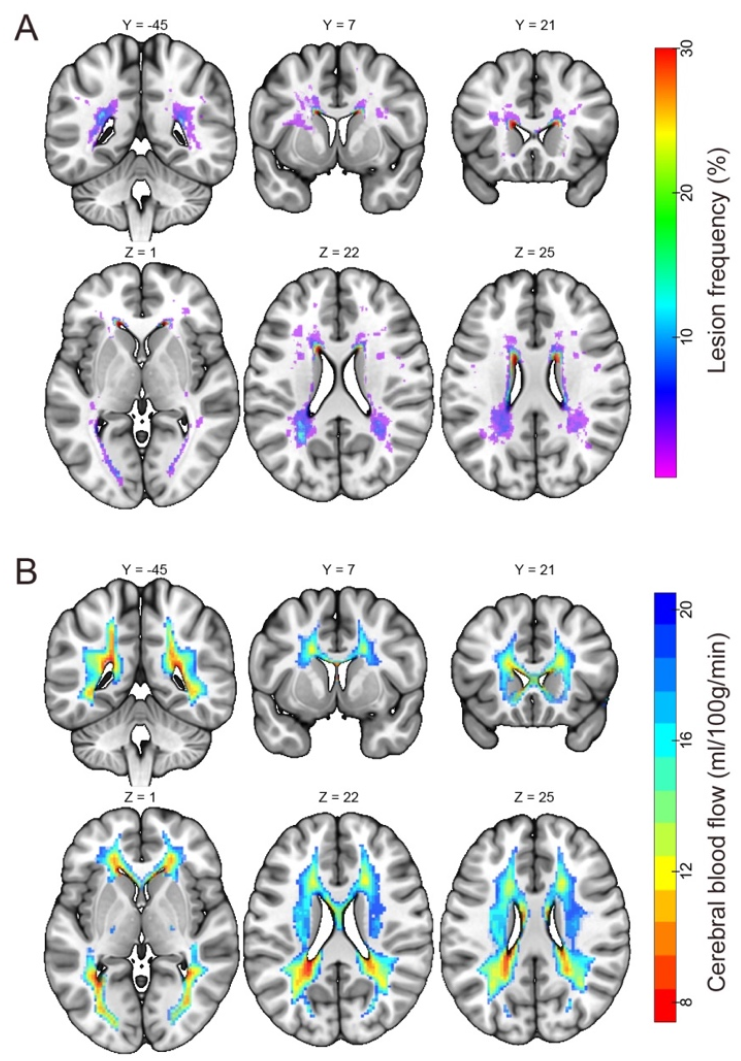
White matter lesion frequency in elderly subjects and cerebral blood flow in middle-aged subjects. (A) Voxel-wise white matter lesion frequency in the elderly cohort (N = 46, shown as percentage), superimposed onto the Montreal Neurological Institute 152 standard space. (B) Group-averaged cerebral blood flow (CBF) map of N = 436 middle aged subjects (28) thresholded to show voxels with CBF in the 0-20 ml/100g/min range, superimposed on the same brain slices as in panel (A).

### Virtual lesion DTI structural disconnectome

Next, we assessed the similarity of DTI disconnectomes caused by WM hyperintensities and pseudo-PVWM lesions simulated based on CBF (Fig. 2). See *Materials and Methods* for further details of these steps. We used the union of the elderly subjects’ WM lesion masks as an ROA and compared it with a virtual PVWM lesion ROA generated using middle-aged (43 – 56 yrs) subjects’ group CBF data with CBF ≤ 16 ml/100g/min. This CBF threshold was chosen to provide a lesion volume that was similar to that observed in the aggregated elderly subjects’ WM lesion masks. HCP DTI data from healthy subjects produced a disconnectome using virtual lesions based on the thresholded CBF masks (Fig. 2B) that was quite similar to that produced by the union mask of the elderly subjects’ WM lesion masks (Fig. 2 A; spatial correlation coefficient r = 0.59, p<10-7; Sorensen-Dice’s similarity coefficient = 0.63) (33, 34). In particular, we observed consistent reductions in many of the connections between parcels, including the subcortex, cerebellum, and visual areas (Fig. 2 A and B). In a post-hoc analysis examining the overlaps of the ROIs affected by the two disconnectomes, we founds extensive overlap in the cerebellum, subcortical, visual system, and default mode network (Fig. 2C). This pattern of results provides preliminary validation for the use of thresholded CBF masks to simulate the disconnectome due to progressive PVWM lesions.

**Figure 2.**
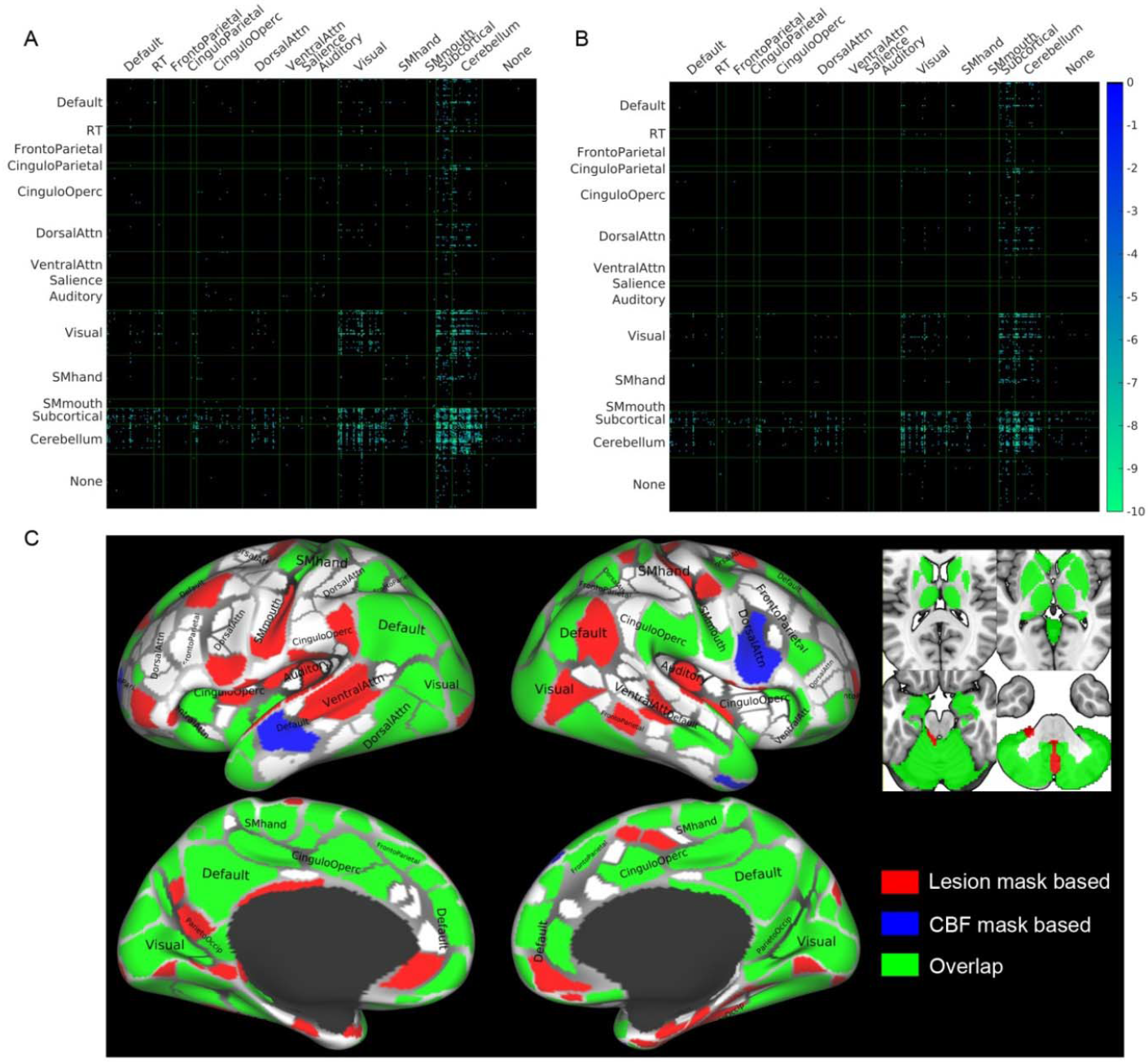
Comparison of disconnectomes using masks based on WM hyperintensities and CBF thresholds. (A, B) Reductions in structural connectivity based on the 30 HCP subjects when using synthesized white matter lesion masks as ROAs in diffusion tractography. The masks used were the union of WM lesion masks of 46 elderly subjects (panel A), and the mask of periventricular white matter CBF ≤ 16 ml/100g/min (panel B), which has similar total lesion volume as that shown in panel A. The matrices show the results of paired t-tests, thresholded at *p* < 0.05 Bonferroni corrected, between structural connectivity with and without an ROA for each of the ROI masks across the 30 subjects. Abbreviation ‘RT’ denotes Retrosplenial Temporal network of the Gordon atlas (36). (C) Overlap of the ROIs affected by the disconnectomes based on the lesion mask in panel A and based on the CBF mask in panel B. For illustration purposes, the cortical parcellations were rendered on the surface using the HCP Connectome WorkBench (37, 38).

To mimic the age-dependent evolution of the WM disconnectome, we next simulated progressive PVWM hyperintensities by varying a threshold applied to the middle aged subjects’ group CBF map (stepwise threshold = 8, 9, 10, …, 19, 20 ml/100g/min as illustrated in Fig. 1B), and used the resulting masks as ROAs for DTI tracking in the 30 HCP subjects’ DTI data. In Fig. 3, we show the network edges of the structural connectome that were significantly reduced in strength (p<0.05, Bonferroni corrected). Connections involving subcortical, cerebellar, and visual regions all showed significant reductions in weight when compared to their homologues in the full connectome. Connections with visual ROIs and some subcortical ROIs were also affected, even with very low CBF masks (see red and orange grid points in Fig. 3).

**Figure 3.**
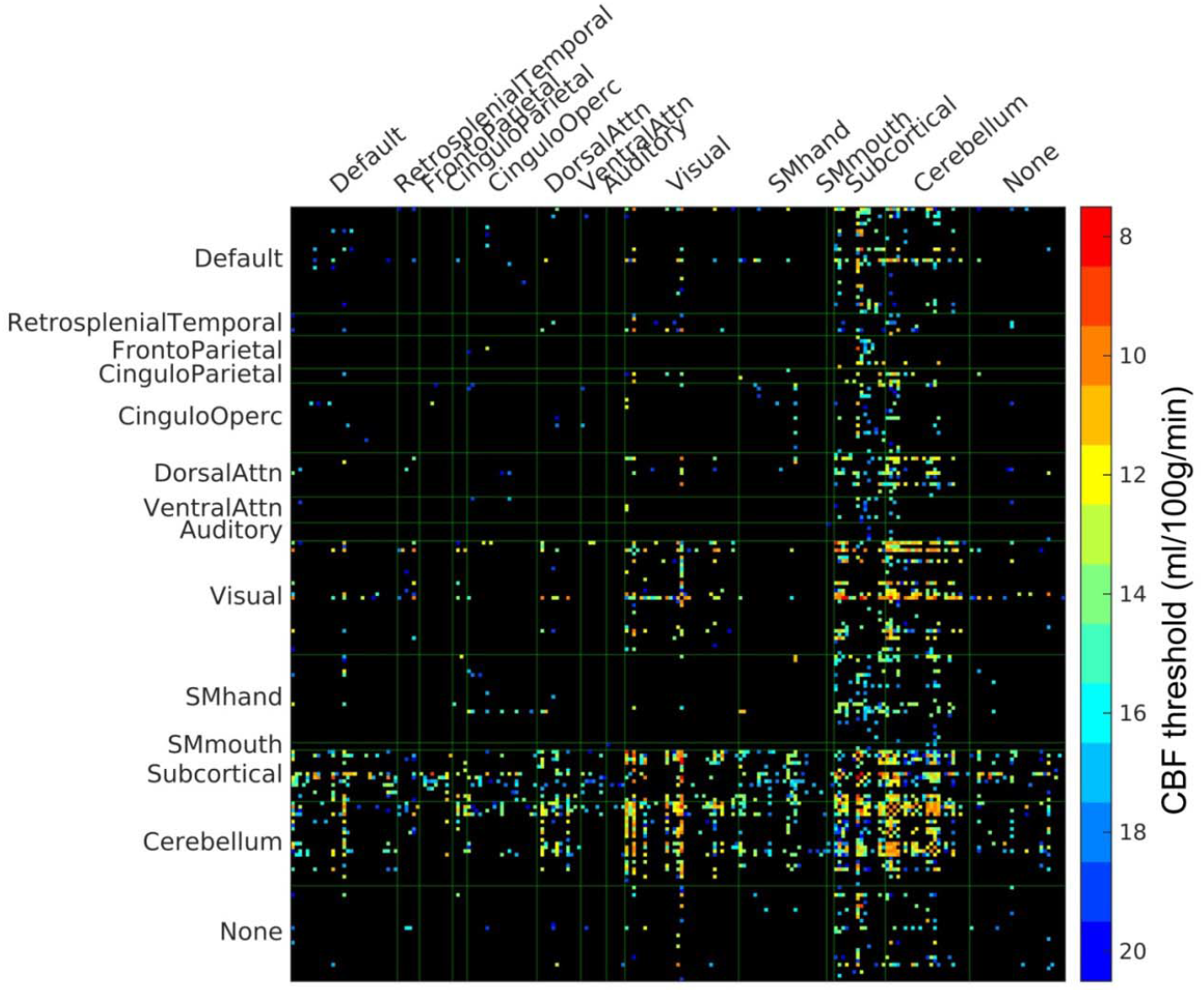
PVWM disconnectome derived using CBF-based progressive PVWM lesion simulation. Edges with significant (*p* < 0.05, Bonferroni corrected) reduction of structural connectome are shown. The color scale shows the ROA mask CBF threshold at which significant disconnection was observed. For illustration purposes, the parcels with no edges of significant change have been eliminated in this matrix, the original full matrix can be found in supplementary Figure S1.

To further elucidate the vulnerability of white matter pathways to progressive PVWM lesion burdens, we conducted streamline tractography of 80 anatomical WM pathways using DSI Studio (35) in the N = 30 healthy HCP subjects and using the step-wise thresholded masks based on the group CBF as the ROAs. The number of affected fibers of each pathway under varying lesion burden was counted and the average percentage of affected fibers was calculated across the 30 subjects. See *Materials and Methods* for further details. In Fig. 4A, we show the average percentage of disconnected streamlines for the affected pathways as a function of simulated progressive PVWM lesion burden, sorted according to lesion burden threshold for 50% disconnection. Optic radiations (OR), inferior fronto-occipital fasciculus (IFOF), inferior longitudinal fasciculus (ILF), corpus callosum (CC), temporopontine tract (TPT), and fornix were the pathways affected under small simulated PVWM lesion burden. Supplementary Fig. S4 shows the results for all 80 pathways. To better illustrate the evolution of affected WM fibers under simulated progressive PVWM lesion burdens, we generated a glass brain view of the affected WM fibers color-coded according to the lesion burden threshold for which they are affected (Fig. 4B) and a time-lapse animation of this evolution in Supplementary Video V1. The glass brain view also demonstrates that OR, IFOF, ILF, CC, and fornix were affected by low PVWM lesion burdens (Fig. 4B).

**Figure 4.**
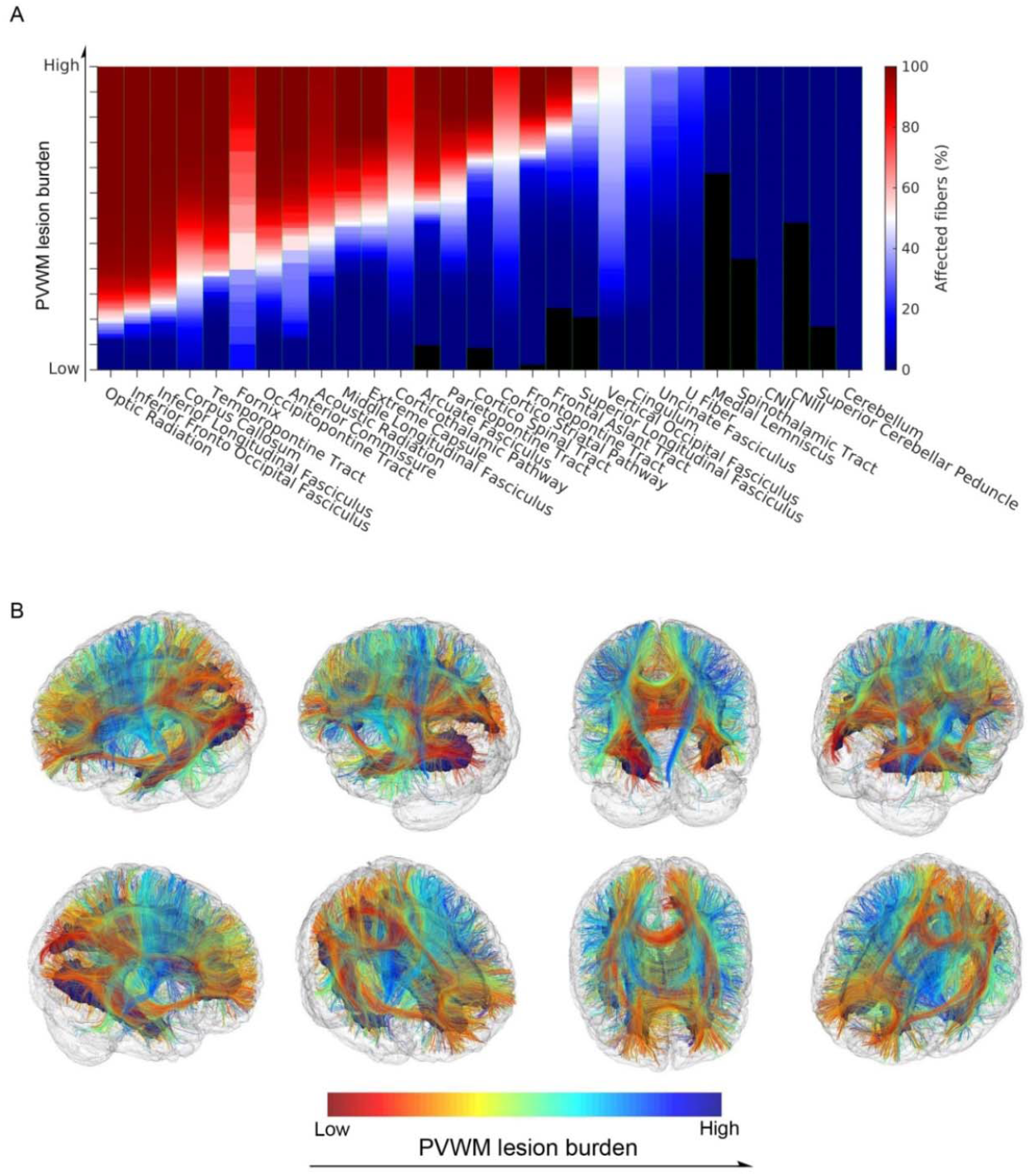
Affected white matter tracts under progressive PVWM lesion burden. Progressive PVWM lesion burdens were simulated by creating virtual lesion masks using a group CBF map stepwise thresholded from low (8 ml/100g/min) to high (20 ml/100g/min) values. (A) Average percentage of affected streamlines of the white matter pathways. The color scale shows the percentage of the number of fibers affected by increasing the simulated lesion burden. The result is averaged across the tractograms of 30 young healthy subjects and across left and right hemispheres. The pathways are ordered by the degree of PVWM burden that affects at least 50% of pathway fibers. (B) Glass brain views of affected white matter fibers with progressive simulated PVWM lesion burden. For visualization purposes, these images were created from the HCP DTI data of a single young healthy subject with streamline fiber tracking (2 x 10^4^ seed points) along with the simulated progressive PVWM lesion as the ROA used to generate the results presented in panel (A). The white matter fibers are color coded to illustrate the onset PVWM lesion burden that affected them.

### Correlations between functional connectivity and DTI connectivity

To obtain additional evidence supporting use of a virtual lesion approach for inferring the effects of WM lesions on structural connectivity, we examined the structure-function relationships in the 46 elderly subjects in whom multimodal MRI data were available, including DTI, resting-state fMRI, and T2-weighted FLAIR MRI (See *Materials and Methods*). Supplementary Figures S2 and S3 show the functional connectivity and structural connectivity matrices, respectively, from a representative elderly subject. The functional connectivity matrix showed high correlations within each of the systems defined by the Gordon functional parcellation (Fig. S2) (36). The structural connectivity matrix was sparser than the functional connectivity matrix, but had some overlapping features (Fig. S3). We compared Spearman’s rank correlations between functional connectivity and different types of structural connectivity in the elderly cohort. The results of these analyses are shown in Figure 5. The strongest correlations between structural and functional connectivity changes were observed when using virtual DTI connectivity (based on the tractography of the healthy HCP subjects’ DTI data) with the elderly subjects’ WM lesion masks as the ROA. Structural connectivity with the subjects’ WM lesion masks as the ROA on their own DTI data showed a higher correlation with functional connectivity than that without adding their WM lesion as an ROA, although the correlation was lower than with connectivity from healthy HCP subjects’ DTI data, even without any ROA. A 3-way ANOVA showed a significant effect of image type (subjects’ real diffusion data or HCP subjects’ diffusion data): *F*(1, 136) = 559.42, *p* < 10-11; and a significant effect of ROA (using the subjects’ WM lesion as ROA or not): *F*(1, 136) = 9.22, *p* < 0.003.

**Figure 5.**
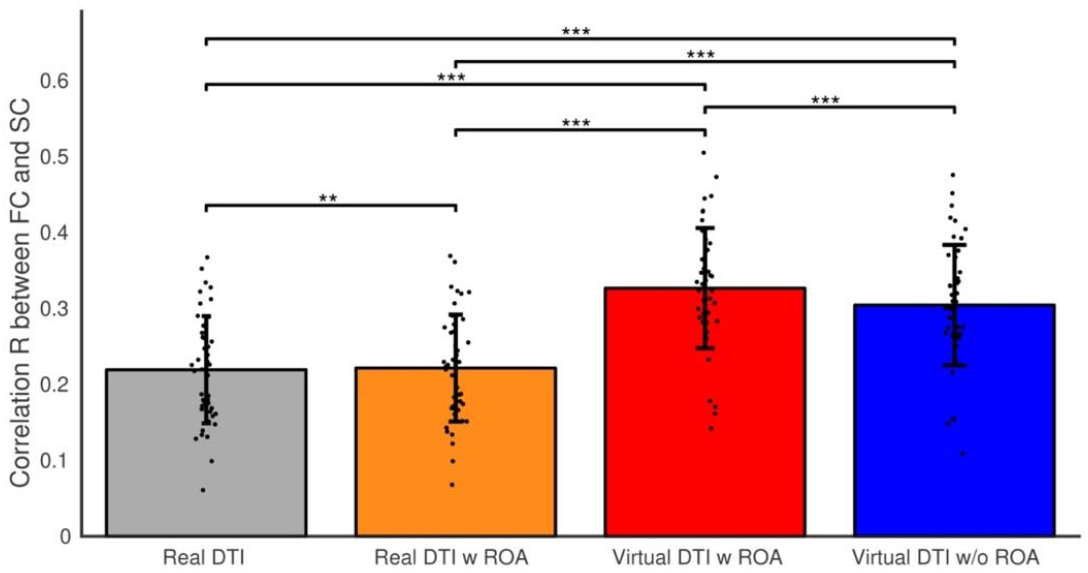
Comparison of FC-SC correlations between real DTI and virtual DTI connectivity. Paired t-tests of functional connectivity (FC) and structural connectivity (SC) correlations between the real DTI connectivity (using elderly subjects’ DTI data) and virtual DTI connectivity (using young adults’ DTI data) with and without using the elderly subjects’ lesion mask as a region of avoidance (ROA). A cross-edges correlation coefficient was calculated for each subject between their FC and SC in the connectivity mask and was compared between the DTI types (real, use the elderly subjects’ own DTI data; virtual, use young HCP subjects’ DTI data; with/without using the subjects white matter lesion as ROA). The significant paired-sample differences are marked with **: *p* < 0.01; ***: *p* < 0.001.

## Discussion

We used a “virtual lesion” approach with healthy subjects’ DTI data from the HCP database and lesion masks as regions of avoidance (ROA) for diffusion tractography to simulate the spatiotemporal changes in structural connectivity associated with progressive ischemic PVWM lesion burden. Use of healthy subject DTI data with virtual lesions circumvented the challenges associated with performing DTI tractography in lesioned brains (24, 25) and progressive ischemic PVWM lesions were simulated by varying the CBF threshold on a mean CBF map from a large cohort of middle-aged subjects, allowing the connectivity effects of PVWM lesions to be isolated from deep WM lesions that are typically seen clinically with advancing age and subcortical ischemic lesion burden. We showed that sub-threshold CBF recapitulates the pattern of PVWM lesion burden seen in an elderly cohort, as demonstrated in our prior work (28).

We validated the virtual lesion approach by assessing correlations between structural and functional connectivity in an elderly cohort in whom multimodal MRI data were available including FLAIR MRI, DTI, and resting-state fMRI, focusing on functional connectivity predicted to be affected by PVWM lesions in the virtual lesion analysis. The structural connectivity of a virtual lesion using the elderly subjects’ WM lesion masks as ROA with young healthy HCP subjects’ DTI data showed a significantly higher correlation with the elderly subjects’ functional connectivity than did the structural connectome using the elderly subjects’ own DTI data. Applying elderly subjects’ WM lesion masks as ROA during tractography on their own DTI data also improved the correlation with their functional connectivity, but to a much lesser extent. This finding was consistent with the notion that the altered diffusion properties in the lesion area can confound fiber tracking algorithms (24, 25). Since structure-function relationships between elderly subject’s functional connectivity data and structural connectivity derived from healthy HCP subjects’ DTI data even without any ROA were also stronger than when using elderly subjects’ DTI data, some of the benefit of the virtual lesion approach seen here is likely also attributable to the higher resolution of HCP DTI data and the increased accuracy (less noise) when using the averaged result of 30 HCP subjects’ DTI scan instead of the elderly subject’s own single DTI scan. Moreover, functional connectivity does not necessarily reflect a single edge; therefore, changes in functional connectivity to compensate for structural connectivity deficits (39–41) may also serve to “normalize” overall functional connectivity. However, to the extent that correlations are expected between structural and functional connectivity, that the strongest correlations were observed when using the virtual lesion DTI approach supports the notion that virtual lesion DTI can be used to infer the effects of WM lesion on structural connectivity.

The virtual lesion approach was used to simulate the effects of progressive ischemic PVWM lesions on both the structural connectome using cortical and subcortical parcellations as seed regions, and on WM pathways defined in DSI Studio. Connectivity to parts of the visual system along with subcortical and cerebellar systems appeared vulnerable to early PVWM lesions, while much more widespread involvement of both cortical and subcortical pathways was seen as simulated PVWM lesion burden increased (see Fig. 4 and Fig S4). Even early PVWM lesions were predicted to affect multiple pathways, likely reflecting the increased density of crossing fibers in the periventricular regions. Pathways including optic radiations, inferior fronto-occipital fasciculus, inferior longitudinal fasciculus, corpus callosum, temporopontine tract and fornix were affected by the smallest simulated PVWM lesion burdens. Early disruption of these pathways is consistent with previous literature suggesting that healthy aging is correlated with reduced integrity of inferior fronto-occipital fasciculus (associated with visual construction) and reduced integrity of inferior longitudinal fasciculus (associated with visualmotor dexterity and visual memory) (42, 43). Aging is also associated with degeneration of the corpus callosum (44, 45) and optic radiations (45), the latter of which typically occur without visual field abnormalities (46). Reduced fornix integrity has also been associated with memory deficits and may predict conversion to cognitive impairment and dementia (47). Partial involvement of multiple pathways by PVWM disease may underlie the reductions in processing speed that characterize age-associated cognitive decline (6).

The use of a virtual lesion approach to identify pathways altered by PVWM is an extension of the “lesion network mapping” approach (48), which uses patients’ traced lesions on healthy subjects’ functional connectivity to identify the connections associated with the patients’ symptoms. Lesion network mapping has thus far been applied primarily to focal lesions such as stroke (49). In the current study, the ability to model the evolution of progressive ischemic PVWM using increasing CBF thresholds to identify vulnerable voxels allows the evolution of network disruption to be characterized. An analogous approach can also be applied to WM lesion frequency maps available from large cohorts (3, 5) as a complementary means of investigating the network consequences of hyperintensities seen on T2-weighted MRI. Correlations between affected pathways and regional cortical atrophy patterns will also contribute to understanding the mechanisms and network consequences of PVWM lesions. Indeed, many of the ROIs affected in the virtual lesion based disconnectome, including calcarine, cuneus, insula, middle and superior frontal, lateral occipital, precentral and angular (Fig. 2C), have been associated with regional cortical atrophy of aging (50).

Since the disconnectome generated here used regions of avoidance (ROA) in DTI tractography, it may overestimate the extent of true disconnection. For example, the data shown in Figure 4A suggest complete or nearly complete disconnection of at least parts of several pathways at higher PVWM lesion burdens, whereas complete deficits such as visual field defects are not typically observed clinically (51). Further, the maximum simulated lesion size of CBF≤ 20 ml/100g/min corresponds to an essentially complete lesion of the PVWM, which rarely occurs clinically. However, the majority of affected pathways do not show complete disconnection, particularly at lower CBF thresholds. There is also mounting evidence that both functional and microstructural changes in the PVWM may precede the development of overt WM lesions based on T2-weighted MRI (20–22). Chronic hypoperfusion observed in the PVWM (7, 52, 53) can alter molecular pathways, disrupting paranodal and axon-glial integrity and resulting in white matter disintegration (54) and impaired signal conduction along affected pathways. Accordingly, it is possible that WM lesions based on T2-weighted MRI alone may underestimate the extent to which WM pathways are affected by progressive PVWM lesions.

Several limitations exist in the presented study. First, the sample size (N = 46) of the elderly subjects was modest. Second, as mentioned above, there are large differences in the diffusion MRI scanning protocols between the HCP subjects and the elderly subjects. Additional, a potential limitation is our choice to use a functionally-defined cortex parcellation combined with an anatomically defined atlas of subcortical and cerebellum, which may favor functional/structural connectivity accordingly. Atlases that are jointly validated using functional and structural connectivity (55) would be beneficial for direct comparisons between functional and structural connectivity. Furthermore, the explored correlation between functional connectivity and structural connectivity assumes that the functional connectivity is exclusively affected by the direct structural connectivity. However, functional connectivity could also be altered by deficits of indirect structural connection (56), state modulation (57), and circadian variations (58). Finally, for computational efficiency, the WM fibers in the current study were extracted using the deterministic fiber tracking method implemented in DSI Studio(59). Compared to probabilistic fiber tracking, deterministic fiber tracking methods are more susceptible to noise and cannot account for the uncertainty in fiber orientation estimation (60). However, a recent study comparing deterministic and probabilistic algorithms suggested that deterministic fiber tracking is well suited for connectome analysis (61). Recent work focusing on white matter voxel graphs for the purposes of connectome based lesion symptom mapping provides an alternative approach for rapidly estimating disconnectomes that could be applied to studies of PVWM lesions (62).

### Conclusions

We used a virtual lesion DTI fiber tracking approach with healthy subject DTI data and a simulated ischemic PVWM lesion mask to estimate the sequence of connectivity changes associated with progressive ischemic PVWM lesions. We found that ischemic PVMW lesions likely cause widespread partial disconnection. The optic radiations, inferior fronto-occipital fasciculus, inferior longitudinal fasciculus, corpus callosum, temporopontine tract and fornix are expected to be affected earliest in progressive PVWM ischemia, while the connectivity of subcortical, cerebellar, and visual regions is expected to be disrupted with increasing simulated PVWM lesion burden. This approach provides a meaningful proxy to the spatial-temporal changes of the brain’s structural connectome under PVWM lesions and can be used to further characterize the cognitive sequelae of PVWM lesions in both normal aging and dementia.

## Materials and Methods

### Subjects

We selected 30 healthy subjects (15 Female, 5 subjects in age range 22-25 yrs, 17 subjects in age range 26-30 yrs, 8 subjects in age range 31-35 yrs) with viable diffusion weighted images from the publicly available Human Connectome Project (HCP) S900 release (63). The subjects’ MRI images were acquired using protocols approved by the Institutional Review Board (IRB) of Washington University in Saint Louis.

We complemented our study of the healthy cohort with multimodal MRI data from 46 cognitively normal elderly subjects (31 female, age = 72.30 ± 6.81 years) from a research study of aging and cognitive impairment conducted at the Penn Memory Center (PMC) and the Alzheimer Disease Core Center (ADCC) at the University of Pennsylvania. All participants had an extensive annual evaluation of medical history, physical examination, neurological history and examination, and psychometric assessment; all of these data are collectively used to determine clinical status (64). The study procedures were approved by the IRB of the University of Pennsylvania. All subjects provided written informed IRB-approved consent prior to participating in the study.

### MRI acquisition

The diffusion MRI images of the HCP subjects were acquired at high resolution (1.25mm isotropic) for connectivity mapping using a customized 3T Siemens scanner. The parameters used were as follows: TR/TE = 5520/89.5ms, 268 x 144 matrix on a 210 x 180 mm FOV, 111 slices. A total of 90 diffusion weighting directions of 3 shells of b = 1000, 2000, and 3000 s/mm^2^ in addition to six b = 0 images were acquired (65).

Elderly subjects recruited at Penn completed an MRI scan using a 3T Siemens Prisma scanner. The scanning protocol included a 0.8mm isotropic T1-MPRAGE structural MRI scan (TR/TE/TI = 2400/2.24/1060ms); an 8-minute multiband BOLD-EPI scan: TR/TE = 720/37ms, flip angle 52°, acquisition matrix = 104 × 104 on a 208 mm x 208 mm FOV, 72 slices with slice thickness = 2 mm, and 420 time points; a 3D 1mm isotropic Flair MRI scan (TR/TE/TI = 6000/289/2200ms), acquisition matrix = 256 x 220, 160 slices; a dual-echo field map scan for field inhomogeneity correction; a diffusion weighted image scan: 1.5mm isotropic (TR/TE = 3027/82.8ms), 140 x 140 matrix on a 120mm x 210mm FOV using multi-shell DTI consisting of three b-values (300, 800, 2000 s/mm^2^) acquired along 15, 30, 64 uniformly distributed directions, respectively, with 9 additional b=0 images acquired. The subjects lay supine in the scanner and small cushions were used to secure the subjects’ heads to minimize head movements during the scans.

### Lesion segmentation

White matter lesions were segmented by the automated lesion growth algorithm (LGA) (66) implemented in the lesion segmentation tool (LST) software (version 2.0.15, https://www.applied-statistics.de/lst.html) for SPM 12. The LGA segments the T1 structural scan into gray and white matter maps, and compares them with the FLAIR intensities to calculate the lesion belief map. The lesion belief map was initially thresholded with the recommended value (cutoff threshold of belief value κ = 0.3) (66) and subsequently grown along voxels with hyper-intensity in the FLAIR image to generate the lesion map.

### Network connectivity analysis

To test the effects of white matter lesions on structural and functional connectivity, we analyzed the network connectivity of the whole brain using the 333 cortex parcellation recently published by Gordon et al. (36), combined with subcortical and cerebellar regions of interest (ROIs) from the AAL atlas (67).

### DTI structural connectivity analysis

DTI structural connectivity was obtained using DSI Studio (59). First the diffusion data were reconstructed using the q-space diffeomorphic reconstruction (QSDR) to calculate the spin orientation distribution of diffusing water in a common stereotaxic space. Then 10^7^ seed points were used for whole brain streamline fiber tracking to generate whole brain WM tracks. The number of tracts connecting each pair of parcellated ROIs were normalized by dividing the sum of the volumes of the paired ROIs (68) to construct the weighted structural connectivity network matrix of the given parcellation. For the network analysis, a group representative DTI connectivity matrix mask was created by retaining edges with the 10% strongest consistency estimates (69), excluding edges between physically nearby ROIs (<20mm) (70).

### Virtual-lesion DTI structural disconnectome

To simulate progressive ischemic PVWM lesions, we created a series of masks using the white matter CBF distribution map from a previous study of a large cohort of middle-aged subjects (28) and by masking that map with thresholds from 8 to 20 ml/100g/min in steps of 1 ml/100g/min. At each step, we used the mask as a region of avoidance (ROA) for DTI structural connectivity tracking on N = 30 DTI scans from young healthy subjects in the HCP S900 release. Paired t-tests were conducted between the connectivity matrices with ROA and the connectivity matrices without ROA to examine the vulnerability of the structural connectivity pattern to the synthetic lesions. Next, we tested the similarity of the disconnectome caused by WM hyperintensities and the disconnectome of PVWM CBF based virtual lesions by examining the spatial correlation and Sorensen-Dice’s similarity coefficient (33, 34) between the two connectomes: (i) virtual-lesion structural connectome using the union of the elderly subjects’ WM hyperintensities masks as an ROA to determine which of the associated structural connections were most vulnerable, (ii) virtual lesion DTI disconnectome of a volume-matched PVWM CBF based lesion. For visualization purposes, we also rendered the cortical and subcortical parcels affected by these two types of disconnectome (37, 38, 71).

To test the vulnerability of the anatomical WM pathways to the progression of ischemic PVWM lesion burden simulated using thresholded CBF masks, we utilized the population-averaged white matter tractography atlas (“HCP842”) published by DSI Studio (version 2019) which contains 80 WM pathways labeled by neuroanatomists to conform to previous neuroanatomical knowledge (35). We conducted streamline tractography (10^7^ seed points) on the 30 HCP subjects to extract the 80 pathways (35, 72) and examined the averaged percentage of the affected fibers of these pathways in the presence of simulated ischemic PVWM lesions created with stepwise CBF thresholds.

### Resting-state functional connectivity analysis

Resting-state fMRI images were first corrected for field inhomogeneity, corrected for motion, coregistered to the T1 image, and warped to the MNI space using Advanced Normalization Tools (ANTs) (73) and custom MATLAB (The Mathworks Inc., Natick, MA) scripts. The motion parameters, white matter, CSF, and global signal (74) were regressed out as nuisance variables from the time series. The mean time series was extracted for each ROI in the parcellation. To assess the brain’s functional network, we estimated the pairwise functional connectivity matrix by calculating the Pearson’s r correlation coefficient between each pair of ROI time series, and then we subsequently transformed the correlation coefficients into z-scores using Fisher’s r-to-z transformation (75).

### Correlation between DTI structural connectivity and resting state functional connectivity

In an effort to further validate a virtual lesion approach for assessing the disconnectome associated with PVWM hyperintensities, we compared the correlations between functional connectivity and 4 types of DTI connectivity: 1) DTI connectivity of the elderly subject’s own real DTI data and their lesion masks as the ROA, 2) DTI connectivity of elderly subject’s own real DTI data without any ROA, 3) the averaged DTI connectivity of 30 HCP healthy subject DTI data with elderly subjects’ lesion masks as ROA, which we will refer to as the virtual lesion DTI, and 4) the averaged DTI connectivity of 30 HCP healthy subject DTI data without any ROA. A network connectivity matrix mask was created by retaining the edges with the 10% strongest consistency estimates in both functional connectivity and DTI connectivity (69), and excluding edges between physically nearby ROIs (<20mm) (70). For each subject, the Spearman’s nonparametric rank correlation coefficient between each type of DTI connectivity and functional connectivity was calculated across the edges in the connectivity mask. A 3-way ANOVA was used to assess the variance of different factors: subject, types of DTI data, and with/without the lesion mask as the ROA. Paired t-tests were conducted to compare these types of DTI connectivity and to assess their similarity to functional connectivity.

## Acknowledgments

This work was supported by NIH grants R01 NS111115, P30 AG010124, P41 EB015893, R03 AG063213 and R01 AG055005. MH was supported in part by The Allen H.and Selma W. Berkman Charitable Trust (Accelerating Research on Vascular Dementia) and NIH (1RF1AG054409 and R01 HL127659-04S1). DB was supported by The Paul G. Allen Family Foundation. The HCP data were provided [in part] by the Human Connectome Project, WU-Minn Consortium (Principal Investigators: David Van Essen and Kamil Ugurbil; 1U54MH091657) funded by the 16 NIH Institutes and Centers that support the NIH Blueprint for Neuroscience Research; and by the McDonnell Center for Systems Neuroscience at Washington University.

## Supplementary Figures

**Supplementary Figure S1.**
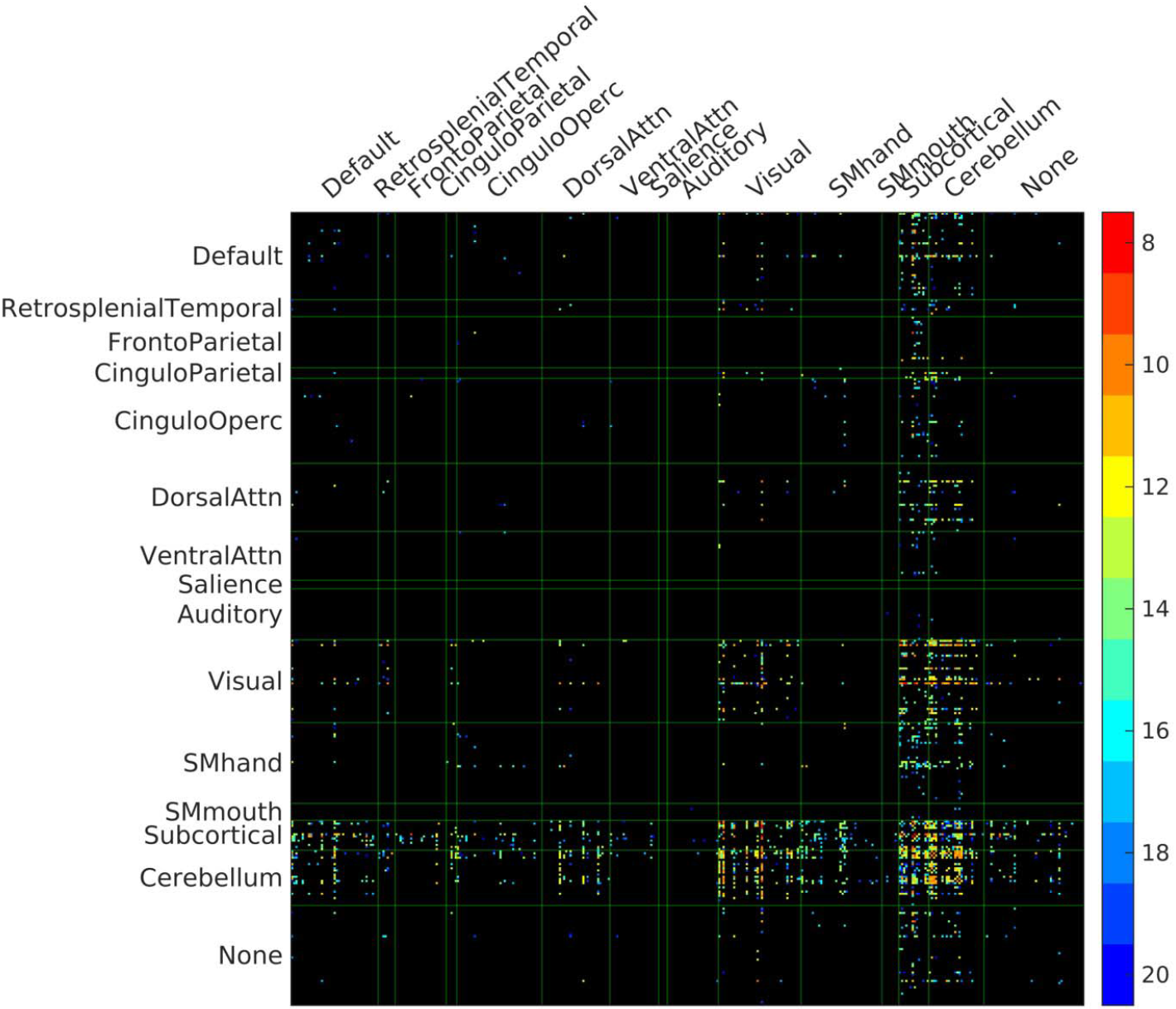
PVWM disconnectome derived using CBF-based progressive PVWM lesion simulation. The connectivity matrix shows edges with significantly reduced structural connectivity (p<0.05, Bonferroni corrected) across N=30 healthy HCP subjects. The color scale shows the ROA mask CBF threshold at which significant disconnection was observed.

**Supplementary Figure S2.**
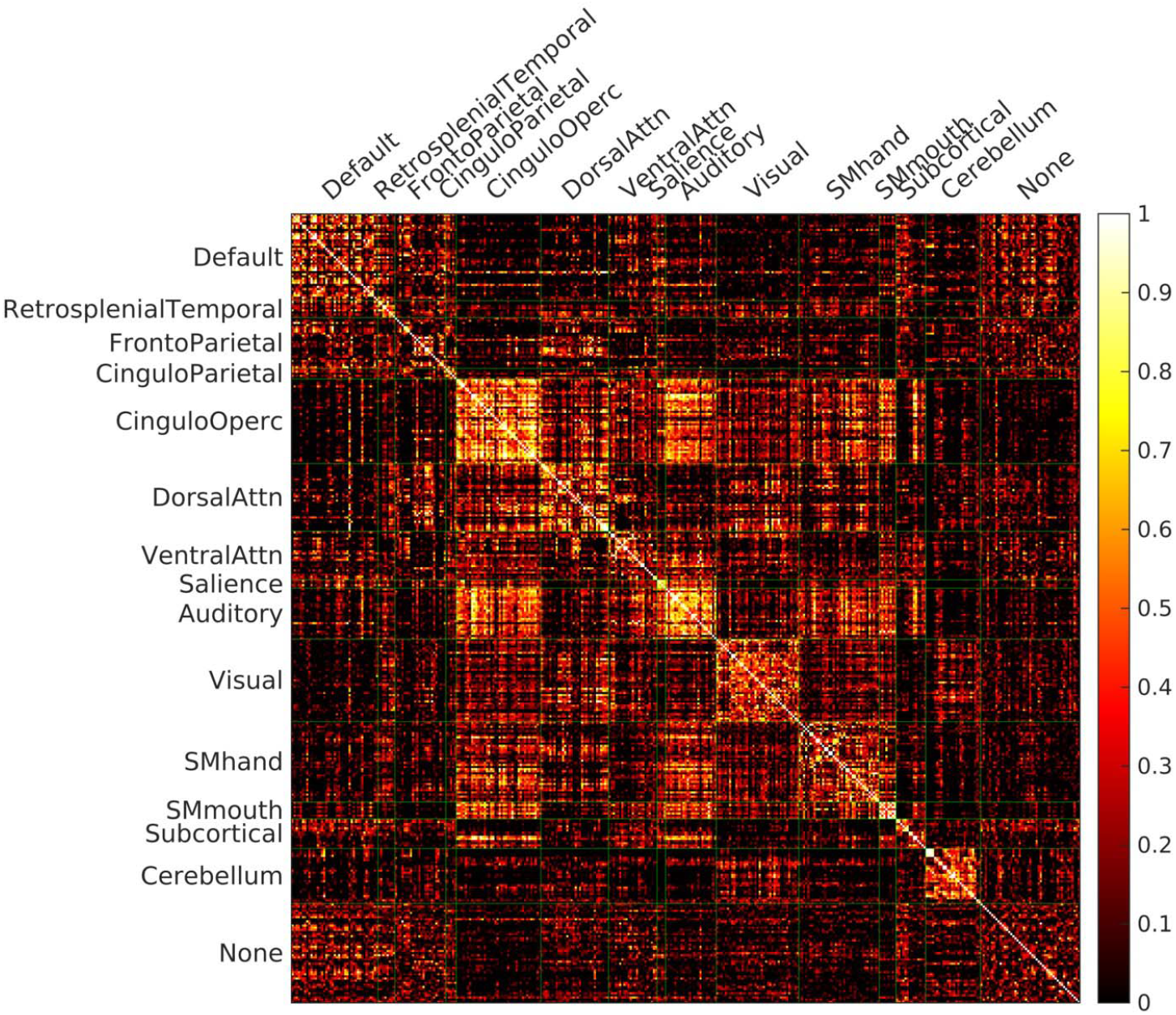
Functional connectivity of an example subject. The functional connectivity was calculated as the correlation coefficient between the time series of pairs of regions. The correlation coefficients were transformed into z-scores using a Fisher r-to-z transformation. For visualization purposes, negative correlations are not shown in this figure.

**Supplementary Figure S3.**
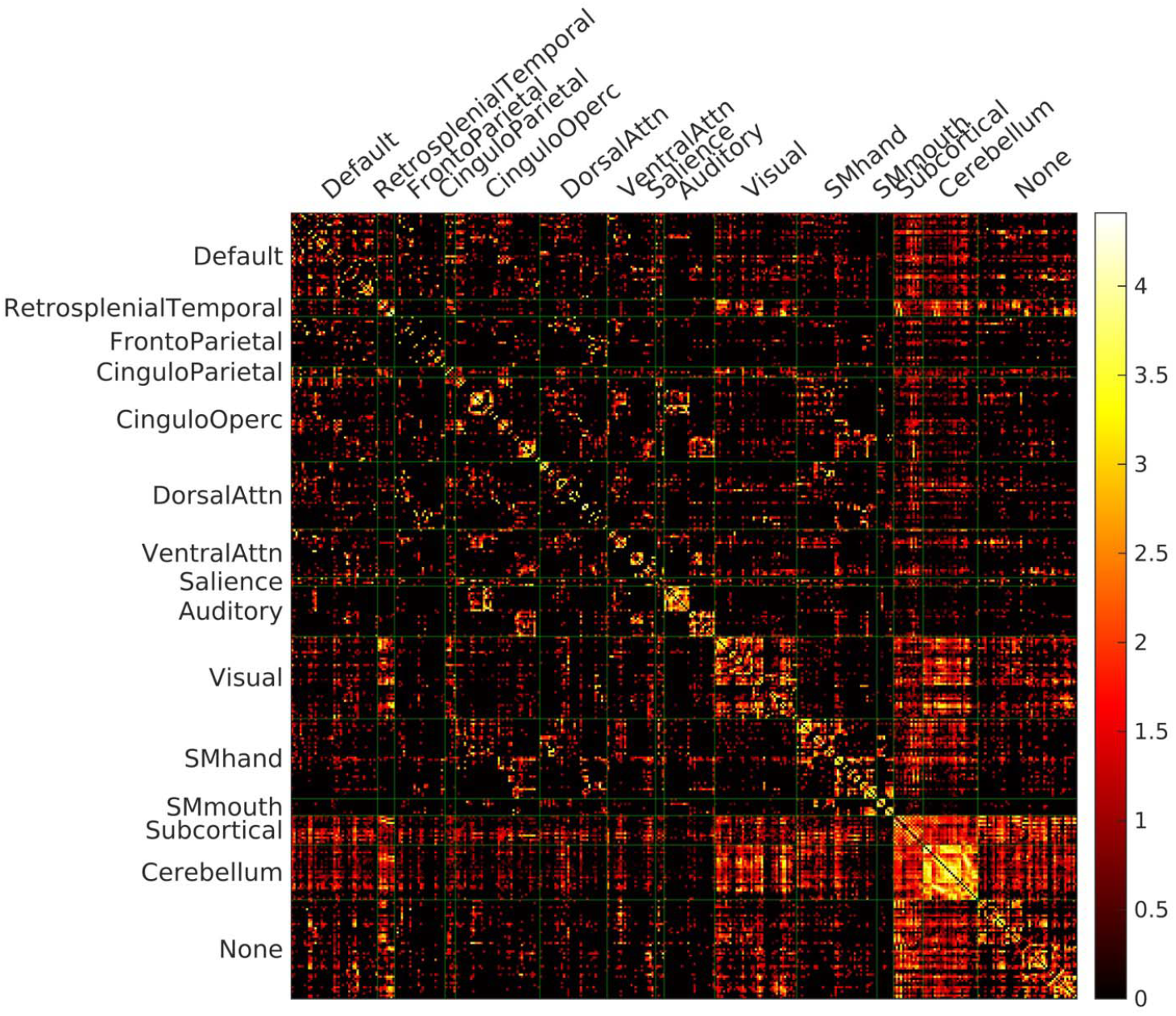
Structure connectivity of an example subject. The number of tracked streamlines were counted between the pairs of regions during DTI fiber tracking, and normalized with the sum of volumes of two ROIs constituting the pair. The structure connectivity is displayed with log10 scale on the right.

**Supplementary Figure S4.**
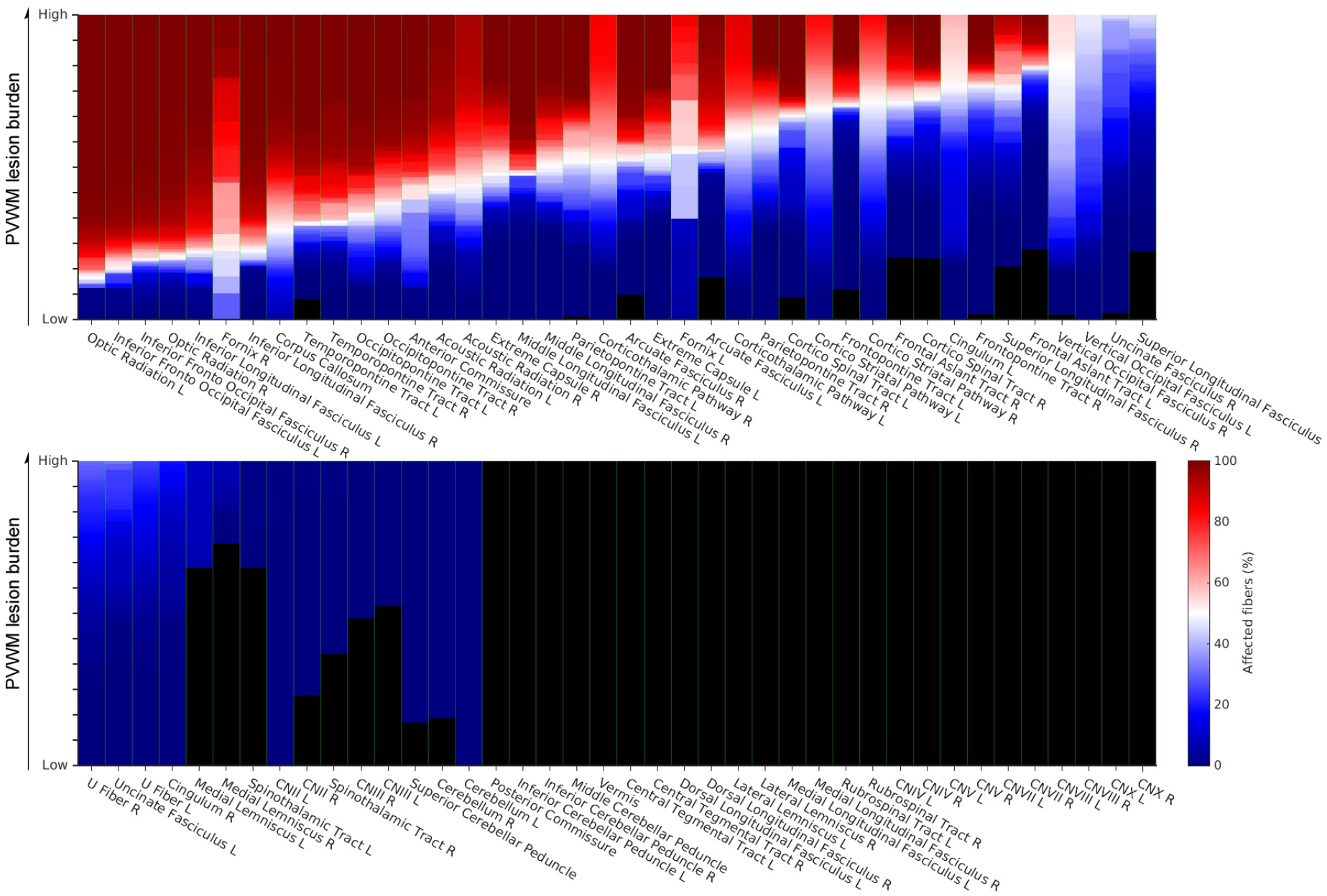
Percentage of affected fibers of the white matter pathways under progressive PVWM lesion burden. The progressive PVWM lesion was simulated by creating lesion masks using a PV CBF map stepwise thresholded at low (8 ml/100g/min) to high (20 ml/100g/min) thresholds. The color scale in the figure encodes the percentage of the number of fibers affected by the simulated lesion burden for each of the 80 pathways. The result is averaged across 30 young healthy subjects’ streamline fiber tracking

## Notes

#### Summary of Updates

1. Fix typos. 2. Removed data availability statement.

## References

1. N. D. Prins, P. Scheltens, White matter hyperintensities, cognitive impairment and dementia: an update. Nat Rev Neurol 11, 157–165 (2015).

2. M. Brant-Zawadzki et al., MR imaging of the aging brain: patchy white-matter lesions and dementia. AJNR Am J Neuroradiol 6, 675–682 (1985).

3. M. Habes et al., White matter lesions: Spatial heterogeneity, links to risk factors, cognition, genetics, and atrophy. Neurology 91, e964–e975 (2018).

4. C. M. Holland et al., Spatial distribution of white-matter hyperintensities in Alzheimer disease, cerebral amyloid angiopathy, and healthy aging. Stroke 39, 1127–1133 (2008).

5. M. Habes et al., White matter hyperintensities and imaging patterns of brain ageing in the general population. Brain 139, 1164–1179 (2016).

6. N. D. Prins et al., Cerebral small-vessel disease and decline in information processing speed, executive function and memory. Brain 128, 2034–2041 (2005).

7. L. Griffanti et al., Classification and characterization of periventricular and deep white matter hyperintensities on MRI: A study in older adults. Neuroimage 170, 174–181 (2018).

8. M. S. Dhamoon et al., Periventricular White Matter Hyperintensities and Functional Decline. J Am Geriatr Soc 66, 113–119 (2018).

9. M. E. Murray et al., A quantitative postmortem MRI design sensitive to white matter hyperintensity differences and their relationship with underlying pathology. J Neuropathol Exp Neurol 71, 1113–1122 (2012).

10. A. A. Gouw et al., Heterogeneity of small vessel disease: a systematic review of MRI and histopathology correlations. J Neurol Neurosurg Psychiatry 82, 126–135 (2011).

11. J. T. O’Brien, H. S. Markus, Vascular risk factors and Alzheimer’s disease. BMC Med 12, 218 (2014).

12. R. N. Kalaria, M. Ihara, Dementia: Vascular and neurodegenerative pathways-will they meet? Nat Rev Neurol 9, 487–488 (2013).

13. A. M. Brickman et al., Reconsidering harbingers of dementia: progression of parietal lobe white matter hyperintensities predicts Alzheimer’s disease incidence. Neurobiol Aging 36, 27–32 (2015).

14. S. C. Larsson, H. S. Markus, Does Treating Vascular Risk Factors Prevent Dementia and Alzheimer’s Disease? A Systematic Review and Meta-Analysis. J Alzheimers Dis 64, 657–668 (2018).

15. E. Bullmore, O. Sporns, Complex brain networks: graph theoretical analysis of structural and functional systems. Nat Rev Neurosci 10, 186–198 (2009).

16. A. J. Lawrence, A. W. Chung, R. G. Morris, H. S. Markus, T. R. Barrick, Structural network efficiency is associated with cognitive impairment in small-vessel disease. Neurology 83, 304–311 (2014).

17. J. Tang et al., Aberrant white matter networks mediate cognitive impairment in patients with silent lacunar infarcts in basal ganglia territory. J Cereb Blood Flow Metab 35, 1426–1434 (2015).

18. A. M. Tuladhar et al., Structural network connectivity and cognition in cerebral small vessel disease. Hum Brain Mapp 37, 300–310 (2016).

19. A. M. Tuladhar et al., Disruption of rich club organisation in cerebral small vessel disease. Hum Brain Mapp 38, 1751–1766 (2017).

20. I. M. Nasrallah et al., White Matter Lesion Penumbra Shows Abnormalities on Structural and Physiologic MRIs in the Coronary Artery Risk Development in Young Adults Cohort. AJNR Am J Neuroradiol 40, 1291–1298 (2019).

21. P. Maillard et al., White matter hyperintensities and their penumbra lie along a continuum of injury in the aging brain. Stroke 45, 1721–1726 (2014).

22. N. Promjunyakul et al., Characterizing the white matter hyperintensity penumbra with cerebral blood flow measures. Neuroimage Clin 8, 224–229 (2015).

23. J. M. Wardlaw, M. C. Valdes Hernandez, S. Munoz-Maniega, What are white matter hyperintensities made of? Relevance to vascular cognitive impairment. J Am Heart Assoc 4, 001140 (2015).

24. W. Reginold et al., Tractography at 3T MRI of Corpus Callosum Tracts Crossing White Matter Hyperintensities. AJNR Am J Neuroradiol 37, 1617–1622 (2016).

25. W. Reginold et al., Impact of white matter hyperintensities on surrounding white matter tracts. Neuroradiology 60, 933–944 (2018).

26. S. Munoz Maniega et al., Spatial Gradient of Microstructural Changes in Normal-Appearing White Matter in Tracts Affected by White Matter Hyperintensities in Older Age. Front Neurol 10, 784 (2019).

27. S. Seiler et al., Cerebral tract integrity relates to white matter hyperintensities, cortex volume, and cognition. Neurobiol Aging 72, 14–22 (2018).

28. S. Dolui et al., Characterizing a perfusion-based periventricular small vessel region of interest. Neuroimage Clin 23, 101897 (2019).

29. C. J. Honey et al., Predicting human resting-state functional connectivity from structural connectivity. Proc Natl Acad Sci U S A 106, 2035–2040 (2009).

30. M. P. van den Heuvel, R. C. Mandl, R. S. Kahn, H. E. Hulshoff Pol, Functionally linked resting-state networks reflect the underlying structural connectivity architecture of the human brain. Hum Brain Mapp 30, 3127–3141 (2009).

31. M. D. Greicius, K. Supekar, V. Menon, R. F. Dougherty, Resting-state functional connectivity reflects structural connectivity in the default mode network. Cereb Cortex 19, 72–78 (2009).

32. M. Straathof, M. R. Sinke, R. M. Dijkhuizen, W. M. Otte, A systematic review on the quantitative relationship between structural and functional network connectivity strength in mammalian brains. J Cereb Blood Flow Metab 39, 189–209 (2019).

33. T. J. Sørensen, A method of establishing groups of equal amplitude in plant sociology based on similarity of species content and its application to analyses of the vegetation on Danish commons (I kommission hos E. Munksgaard, 1948).

34. L. R. Dice, Measures of the amount of ecologic association between species. Ecology 26, 297–302 (1945).

35. F. C. Yeh et al., Population-averaged atlas of the macroscale human structural connectome and its network topology. Neuroimage 178, 57–68 (2018).

36. E. M. Gordon et al., Generation and Evaluation of a Cortical Area Parcellation from Resting-State Correlations. Cereb Cortex 26, 288–303 (2016).

37. D. S. Marcus et al., Informatics and data mining tools and strategies for the human connectome project. Front Neuroinform 5, 4 (2011).

38. D. C. Van Essen, M. F. Glasser, D. L. Dierker, J. Harwell, T. Coalson, Parcellations and hemispheric asymmetries of human cerebral cortex analyzed on surface-based atlases. Cereb Cortex 22, 2241–2262 (2012).

39. D. Barulli, Y. Stern, Efficiency, capacity, compensation, maintenance, plasticity: emerging concepts in cognitive reserve. Trends Cogn Sci 17, 502–509 (2013).

40. A. Celeghin et al., Intact hemisphere and corpus callosum compensate for visuomotor functions after early visual cortex damage. Proc Natl Acad Sci U S A 114, E10475–E10483 (2017).

41. A. Iraji et al., Compensation through Functional Hyperconnectivity: A Longitudinal Connectome Assessment of Mild Traumatic Brain Injury. Neural Plast 2016, 4072402 (2016).

42. A. N. Voineskos et al., Age-related decline in white matter tract integrity and cognitive performance: a DTI tractography and structural equation modeling study. Neurobiol Aging 33, 21–34 (2012).

43. N. Shinoura et al., Impairment of inferior longitudinal fasciculus plays a role in visual memory disturbance. Neurocase 13, 127–130 (2007).

44. M. Ota et al., Age-related degeneration of corpus callosum measured with diffusion tensor imaging. Neuroimage 31, 1445–1452 (2006).

45. A. Peters, The effects of normal aging on myelin and nerve fibers: a review. J Neurocytol 31, 581–593 (2002).

46. M. Kitajima et al., Hyperintensities of the optic radiation on T2-weighted MR images of elderly subjects. AJNR Am J Neuroradiol 20, 1009–1014 (1999).

47. V. Douet, L. Chang, Fornix as an imaging marker for episodic memory deficits in healthy aging and in various neurological disorders. Front Aging Neurosci 6, 343 (2014).

48. R. R. Darby, J. Joutsa, M. J. Burke, M. D. Fox, Lesion network localization of free will. Proc Natl Acad Sci U S A 115, 10792–10797 (2018).

49. M. D. Fox, Mapping Symptoms to Brain Networks with the Human Connectome. N Engl J Med 379, 2237–2245 (2018).

50. A. Bakkour, J. C. Morris, D. A. Wolk, B. C. Dickerson, The effects of aging and Alzheimer’s disease on cerebral cortical anatomy: specificity and differential relationships with cognition. Neuroimage 76, 332–344 (2013).

51. D. Pascolini, S. P. Mariotti, Global estimates of visual impairment: 2010. Br J Ophthalmol 96, 614–618 (2012).

52. L. Pantoni, J. H. Garcia, Pathogenesis of leukoaraiosis: a review. Stroke 28, 652–659 (1997).

53. K. W. Kim, J. R. MacFall, M. E. Payne, Classification of white matter lesions on magnetic resonance imaging in elderly persons. Biol Psychiatry 64, 273–280 (2008).

54. M. M. Reimer et al., Rapid disruption of axon-glial integrity in response to mild cerebral hypoperfusion. J Neurosci 31, 18185–18194 (2011).

55. L. Fan et al., The Human Brainnetome Atlas: A New Brain Atlas Based on Connectional Architecture. Cereb Cortex 26, 3508–3526 (2016).

56. C. D. Langen et al., White matter lesions relate to tract-specific reductions in functional connectivity. Neurobiol Aging 51, 97–103 (2017).

57. Z. Li et al., Effects of resting state condition on reliability, trait specificity, and network connectivity of brain function measured with arterial spin labeled perfusion MRI. Neuroimage 173, 165–175 (2018).

58. B. J. Shannon et al., Morning-evening variation in human brain metabolism and memory circuits. J Neurophysiol 109, 1444–1456 (2013).

59. F. C. Yeh, V. J. Wedeen, W. Y. Tseng, Estimation of fiber orientation and spin density distribution by diffusion deconvolution. Neuroimage 55, 1054–1062 (2011).

60. D. K. Jones, Studying connections in the living human brain with diffusion MRI. Cortex 44, 936–952 (2008).

61. T. Sarwar, K. Ramamohanarao, A. Zalesky, Mapping connectomes with diffusion MRI: deterministic or probabilistic tractography? Magn Reson Med 81, 1368–1384 (2019).

62. C. Greene et al., Finding maximally disconnected subnetworks with shortest path tractography. Neuroimage Clin 23, 101903 (2019).

63. D. C. Van Essen et al., The WU-Minn Human Connectome Project: an overview. Neuroimage 80, 62–79 (2013).

64. S. Dolui et al., Comparison of PASL, PCASL, and background-suppressed 3D PCASL in mild cognitive impairment. Hum Brain Mapp 38, 5260–5273 (2017).

65. S. N. Sotiropoulos et al., Advances in diffusion MRI acquisition and processing in the Human Connectome Project. Neuroimage 80, 125–143 (2013).

66. P. Schmidt et al., An automated tool for detection of FLAIR-hyperintense white-matter lesions in Multiple Sclerosis. Neuroimage 59, 3774–3783 (2012).

67. E. Mellet et al., Neural basis of mental scanning of a topographic representation built from a text. Cereb Cortex 12, 1322–1330 (2002).

68. E. van Dellen et al., Minimum spanning tree analysis of the human connectome. Hum Brain Mapp 39, 2455–2471 (2018).

69. J. A. Roberts, A. Perry, G. Roberts, P. B. Mitchell, M. Breakspear, Consistency-based thresholding of the human connectome. Neuroimage 145, 118–129 (2017).

70. J. D. Power et al., Functional network organization of the human brain. Neuron 72, 665–678 (2011).

71. D. C. Van Essen et al., The Brain Analysis Library of Spatial maps and Atlases (BALSA) database. Neuroimage 144, 270–274 (2017).

72. F. C. Yeh, W. Y. Tseng, NTU-90: a high angular resolution brain atlas constructed by q-space diffeomorphic reconstruction. Neuroimage 58, 91–99 (2011).

73. B. B. Avants et al., A reproducible evaluation of ANTs similarity metric performance in brain image registration. Neuroimage 54, 2033–2044 (2011).

74. M. D. Fox, D. Zhang, A. Z. Snyder, M. E. Raichle, The global signal and observed anticorrelated resting state brain networks. J Neurophysiol 101, 3270–3283 (2009).

75. R. A. Fisher, On the probable error of a coefficient of correlation deduced from a small sample. Metron 1, 3–32 (1921).

